# *Candida* Species Isolated in Female Patients of Reproductive Age with Vaginal Candidiasis in Gualeguaychú, Entre Ríos, Argentina

**DOI:** 10.1101/2024.11.02.620544

**Authors:** Pérez Duarte Iván Rodrigo, Razetto Georgina, Leiva Silvina Érica, Torres Luciano, Juárez María Josefina

**Affiliations:** Biochemist, Microbiology Laboratory, Gualeguaychú Centenary hospital, Gualeguaychú, Entre Ríos, Argentina; Graduate Member of the Microbiology Laboratory, Gualeguaychú Centenary hospital, Gualeguaychú, Entre Ríos, Argentina; Biochemist Specialist in Bacteriology, Member of the Microbiology Laboratory, Gualeguaychú Centenary hospital, Gualeguaychú, Entre Ríos, Argentina. Head of the Microbiology Area, Gualeguaychú Centenary hospital, Gualeguaychú, Entre Ríos, Argentina. Microbiology Laboratory, Gualeguaychú Centenary hospital, Gualeguaychú, Entre Ríos, Argentina

**Keywords:** Vaginal Candidiasis, *Candida*, Susceptibly, Antifungal

## Abstract

*Objective*: To identify the *Candida* species isolated in female patients of reproductive age with vaginal candidiasis. To determine the antifungal (ATF) sensitivity profile of the isolated *Candida* species. *Materials and methods*: Cross-sectional descriptive study, in which 124 *Candida spp* strains from vaginal discharge samples were isolated in Sabouraud medium supplemented with glucose. *CHRO Magar Candida* was used for species identification and complementary tests such as germ tube, chlamydoconidia development and investigation of trehalose assimilation were performed. *In vitro* sensitivity was investigated by diffusion method with ATF discs fluconazole (FLU), itraconazole (ITRA) and amphotericin B (AMB). *Results: C. albicans* was observed in 85.5% of the isolates followed by *C. glabrata* with 7.3%, *C. krusei* 4%, *C. tropicalis* 2.4% and other *Candida* species. For FLU, 0.9% of *C. albicans*, 11.1% of *C. glabrata* and 100% of *C. krusei* isolates showed resistance. For ITRA, 17% of *C. albicans* isolates, 55.6% of *C. glabrata* and 100% of *C. krusei* were resistant. There were no isolates resistant to AMB. *Conclusions*: Vaginal candidiasis continues to present *Candida albicans* as the main etiological agent, which is widely sensitive to ATFs. *C. glabrata* and *C. krusei* species show increased resistance to azoles. The results obtained ratify the growing need for *Candida* species identification tests and determination of *in vitro* sensitivity to ATFs in order to guide the treatment of vaginal candidiasis towards therapeutic success.

## 1. Introduction

Vaginal candidiasis in women of reproductive age is a frequent cause of gynecological consultation. Approximately 75% of women will develop this infection at least once in their lifetime. Although a small percentage develop recurrent episodes (at least four episodes per year), it is of utmost importance to achieve a correct antifungal treatment in order to improve the quality of life of women who suffer from it [2, 4, 5]

*Candida* yeasts present in the vagina are part of the normal microbiota, which is considered an aspect that can cause complications when diagnosing this condition, because it is unknown when the amount of yeast present in the vagina begins to produce pathological effects in the vagina [2].

Vaginal *Candida* infections are usually of endogenous origin due to modification of the vaginal microbial flora, either after antibiotic treatment or as a result of a decrease in the host’s immune response as in the case of treatment with corticosteroids, uncontrolled diabetes, immune-compromised diseases, obesity, stress or use of contraceptives [4, 5].

The etiological agent mostly isolated is *Candida albicans*, widely sensitive to antifungals [1, 3, 10, 17]. The persistence or recurrence of infections is usually attributed to *non-albicans Candida* species, such as *C. glabrata, C. krusei* as causative agents of vaginal pathology. These species are usually the most associated with episodes of recurrence because they are more resistant to azoles, such as FLU and ITRA [10, 11, 20].

## 2. Materials and Methods

With prior authorization, analysis and approval of the work protocol by the Head of the Central Laboratory and the Teaching and Research Committee of the Gualeguaychú Centenary Hospital, the storage process of *Candida spp* strains obtained from vaginal discharge samples of women of reproductive age with signs and symptoms compatible with candidiasis at the vaginal level was developed between May 2022 and January 2023 at the Microbiology Laboratory of the Gualeguaychú Centenary Hospital.

Identification of clinical isolates:

The different samples of vaginal discharge were processed by direct mycological examination and cultures in different media. For the identification of the genus *Candida*, the macro and microscopic morphology of the colonies, presence or absence of capsule, pigment production, size, shape and growth at differential temperatures were taken into account.

First, the yeasts were isolated on Sabouraud agar medium supplemented with glucose (400 mg/dL) and chloramphenicol (5 mg/dL) (Britania brand), incubated at 37°C under aerobic conditions, with final reading between 24 and 48 hours, selecting those belonging to the genus *Candida*. To begin the process of identification of *Candida* species, we proceeded to inoculate *plates of CHROMagar Candida (CHROMagarTM Candida, registered trademark Dr. A Rambach, Paris, France)*, from the cultures developed on Sabouraud glucose-chloramphenicol agar. This is a selective and orienting chromogenic agar medium for the development of *Candida* species complexes that allows differentiation of colonies according to the color and morphology developed [9].

The culture conditions in this selective and differential medium require incubation at 37°C, in an aerobic manner, with final reading between 24 and 48 hours, being useful to guide the typing of the strains under study, which will later be confirmed and classified into species by performing complementary tests such as germ tube formation in human serum pool and chlamydoconidia development in cornmeal-tween 80 agar medium to identify *Candida albicans/Candida dubliniensis* complex, trehalose assimilation research for *Candida glabrata* complex, pseudohyphae formation in human serum pool (*C. tropicalis*), *and* for *C. krusei*, pseudohyphae formation in human serum pool (*C. tropicalis*). *tropicalis*) *and* as for *C. krusei*, the hydrolysis of the Urea molecule was used as an identification test, which in this particular study all the strains of *C. krusei* gave positive results, while the rest of the Candida species gave negative results to this test [9, 26, 27, 28].

Determination of *in vitro* sensitivity:

From pure yeast colonies already identified in genus and species, a 0.5 McFarland turbidity inoculum was prepared in 0.15 M NaCl solution (0.85% saline).

For the diffusion method, 10 cm diameter Petri dishes were used, with 25 ml of Mueller-Hinton agar (Britania), supplemented with 2% glucose and methylene blue, in a final concentration of 0.5 mg/ml, receiving the name of modified Mueller-Hinton.

Using sterile swabs previously dipped in the 0.5 McFarland turbidity inocula of each strain under study, the surface of the plates was seeded with modified Mueller-Hinton in two directions. ATF FLU (25 ug), ITRA (10 ug) and AMB (10 ug) discs (Neo-Sensitabs®; Rosco Diagnóstika) were then plated and incubated at 37°C for 24-48 hours. After this time, the diameter of the growth inhibition zone expressed in millimeters (mm) was measured and classified as “Sensitive” or “Resistant”. Particularly, the *C. glabrata-Fluconazole* combination was classified as ‘‘Sensitive Dose Dependent (SDD) or ‘‘Resistant’’. The cut-off points for FLU, ITRA and AMB were those established by the manufacturer, Neo-Sensitabs®; Rosco Diagnóstika [13, 29].

## 3. Result

A total of 124 strains of *Candida spp* (n=124), corresponding to vaginal discharge samples from women of childbearing age, submitted to the Microbiology Laboratory of the Centennial Hospital of Gualeguaychú, were analyzed. It was determined that 85.5% of the isolates corresponded to *Candida albicans* strains, followed by *C. glabrata* 7.3%, *C. krusei* 4.0%, *C. tropicalis* 2.4% and finally 0.8% for other species (Table 1) and (Figure 1).

**Table 1.**
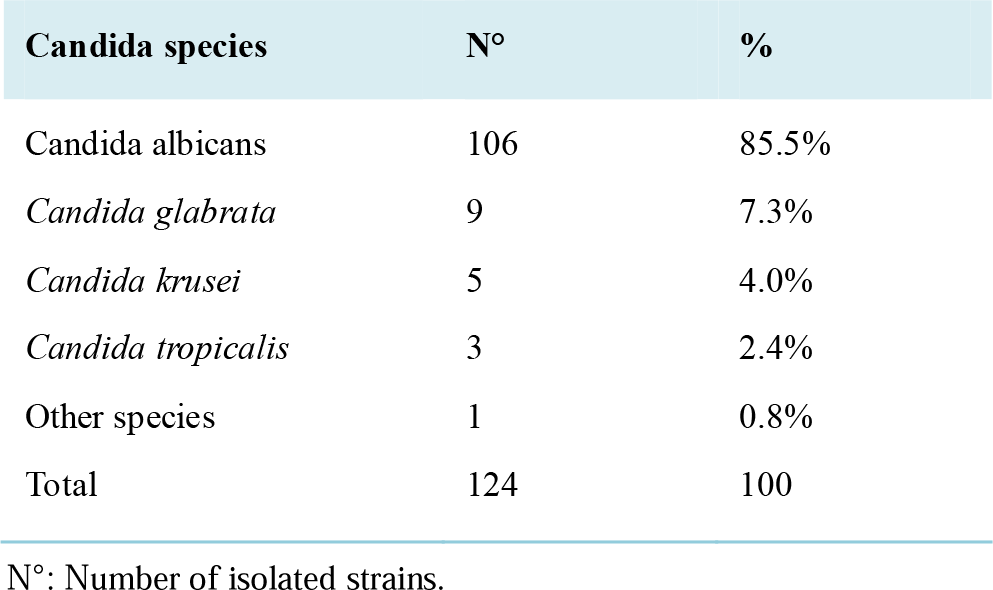
Percentage distribution of Candida species isolated from vaginal discharge samples.

| **Candida species** | **N°** | **%** |
| --- | --- | --- |
| *Candida albicans* | 106 | 85.5% |
| *Candida glabrata* | 9 | 7.3% |
| *Candida krusei* | 5 | 4.0% |
| *Candida tropicalis* | 3 | 2.4% |
| Other species | 1 | 0.8% |
| Total | 124 | 100 |

Nº: Number of isolated strains.

**Figure 1.**
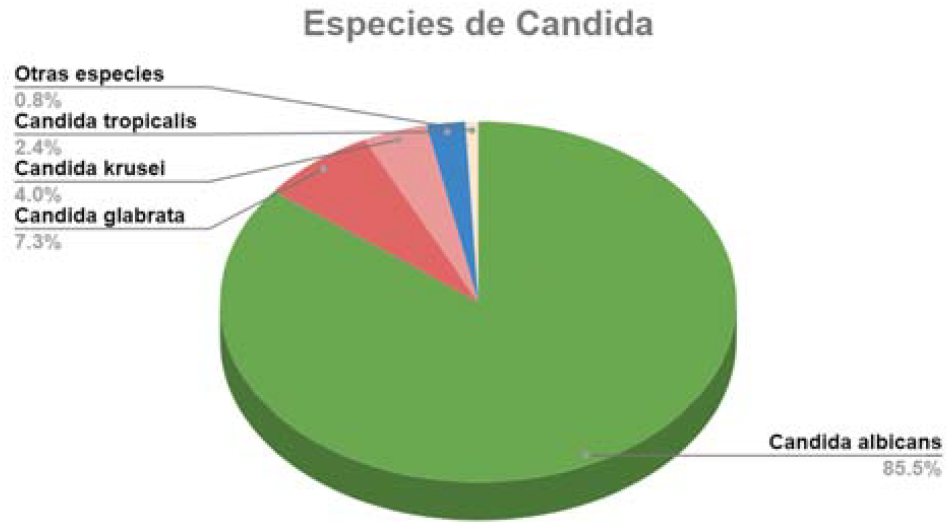
Percentage graph of Candida species isolated.

Regarding susceptibility to FLU, Figure 2 shows that 0.9% of *Candida albicans* isolates were resistant, while 11.1% *of Candida glabrata* isolates were resistant (the remaining 88.9% were dose-dependent sensitive) and no resistance was observed for *Candida tropicalis* isolates. The 5 isolates of *Candida krusei* were resistant to FLU, which is a species with intrinsic resistance to this antifungal agent, through the azole-induced increase in the expression of membrane transporter pumps [24].

**Figure 2.**
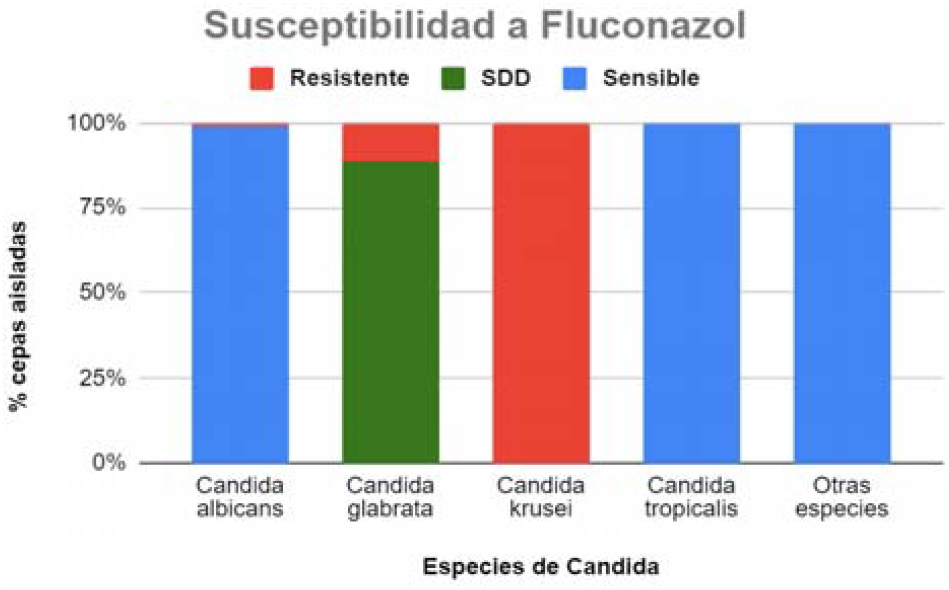
Distribution of susceptibility to Fluconazole of the isolated Candida species. Dose-dependent sensitivity of C. glabrata. Intrinsic resistance of C. krusei.

Figure 3 shows the susceptibility to ITRA, in which it is evident that 17.0% of the isolates of *C. albicans* presented resistance, with respect to the isolates of *C. glabrata* these presented 55.6% resistance, the isolates of *C. tropicalis* and other species did not present resistance. *C. krusei* isolates showed 100% resistance to ITRA.

**Figure 3.**
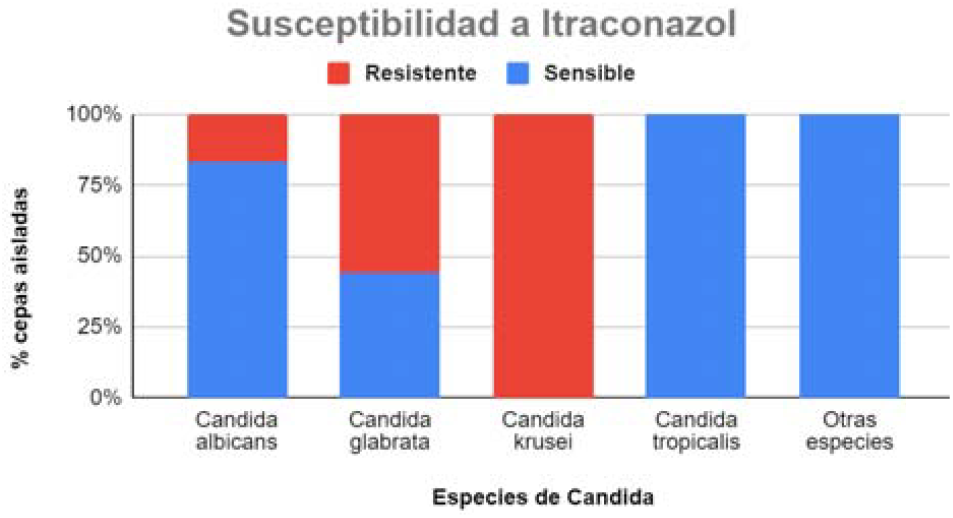
Itraconazole susceptibility distribution of isolated Candida species.

Figure 4 shows that the isolates of *C. albicans, C. glabrata, C. krusei, C. tropicalis* and other species did not show resistance, i. e. they were 100% sensitive to AMB. Although AMB has the best sensitivity profile of the three TFAs, it is not considered the antifungal of choice for vaginal candidiasis, but it is used as one of the best options for the treatment of deep mycosis [5, 21].

**Figure 4.**
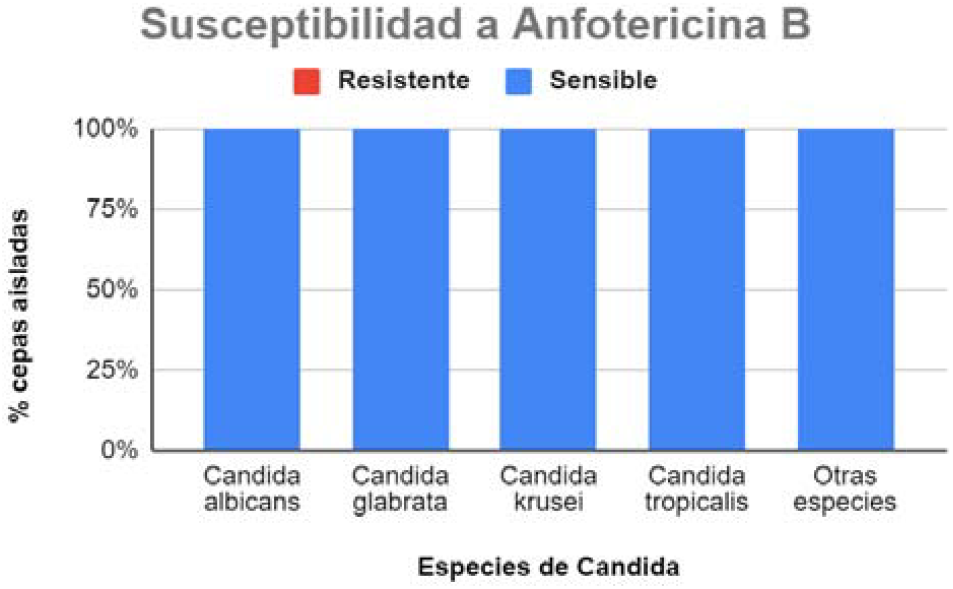
Distribution of susceptibility to amphotericin B of the isolated Candida species.

## 4. Discussion

Vaginal candidiasis is a very common infection in women of reproductive age that usually causes discomfort such as itching, irritation, burning, secretions, slightly unpleasant odor and in extreme cases can even alter the patient’s behavior, causing depression, moodiness and anxiety. It is a common and uncomplicated pathology within the public health system. It is not advisable to offer empirical treatments, based on the clinic, in those vaginitis considered mild, moderate or complex, where there is a tendency to use azoles (FLU, ITRA) in oral form or by means of ovules.) For all the above mentioned situations, microbiological identification of the yeast and its susceptibility to ATF [6] is recommended.

Studies carried out in recent years show that *C. albicans* is the species most responsible for vaginal candidiasis in women of reproductive age (75% to 95%), while other species, *C. glabrata* and *C. krusei*, are found less frequently [7, 8, 12, 14, 15]. In the present study, like the results obtained in the selected literature, *C. albicans* was the most frequently isolated species, followed by *C. glabrata, C. krusei* and *C. tropicalis*. The results of the susceptibility tests reflect that resistance to ATF was low for the genus *Candida*, which is in agreement with research conducted by Dalben *et al*. [21], Richter *et al*. [25] and other authors [5, 11, 22, 23, 24]. However, it is possible that due to empirically indicated treatments, low adherence to FLU and ITRA treatments, or in cases of women who resort to self-medication (as a consequence of the easy availability of these drugs), this is contributing to the increase in the population of ATF-resistant yeasts [6]. The increase in the resistance of *non-albicans Candida* species such as *C. krusei, C. glabrata*, which are usually associated with failures in empirically applied treatments, can be highlighted [5, 16, 18, 19]. It is important to point out that the identification of all strains causing vaginal candidiasis together with the corresponding ATF susceptibility tests are favorable for an accurate choice of antifungal therapy and to ensure a favorable clinical evolution of the patients.

## 5. Conclusions

The present work was a descriptive study whose objective was to characterize vaginitis caused by *Candida* in the city of Gualeguaychú, using data obtained from the Microbiology Laboratory of the Gualeguaychú Centenary Hospital. However, it should be considered that these are elementary epidemiological results; in order to be closer to reality, it is necessary to resort to a larger number of data obtained by official organizations. From the information obtained it can be inferred: 1) as in the rest of Argentina and other Latin American countries, vaginal candidiasis still has *Candida albicans* as the main etiological agent, 2) it can be concluded that, in the studied population of women of fertile age with candidiasis vaginitis, *Candida* yeasts are still widely sensitive to antifungals, 3) the species *C. glabrata and C*. 4) The results obtained with strains isolated from vaginal discharge samples confirm the growing need to carry out tests for identification of *Candida* species and determination of *in vitro* sensitivity to antifungals in order to guide the treatment of vaginal candidiasis towards therapeutic success and avoid recurrent or chronic infections.

## Abbreviations

ATF: Antifungal
FLU: Fluconazole
ITRA: Itraconazole
AMB: Amphotericin B

## Acknowledgments

Head laboratory Bioq. Siri, Leticia; Lic. Urriste, Celeste; Dr. Bioq. Levin, Gustavo.

